# MARQO pipeline resolves multiparametric cellular and spatial organization in cancer tissue lesions

**DOI:** 10.1101/2025.04.24.650539

**Authors:** Mark Buckup, Igor Figueiredo, Giorgio Ioannou, Sinem Ozbey, Rafael Cabal, Alexandra Tabachnikova, Leanna Troncoso, Jessica Le Berichel, Zhen Zhao, Stephen C. Ward, Clotilde Hennequin, Guray Akturk, Steve Hamel, Maria Isabel Fiel, Rachel Brody, Myron Schwartz, Thomas U. Marron, Seunghee Kim-Schulze, Edgar Gonzalez-Kozlova, Vladimir Roudko, Pauline Hamon, Miriam Merad, Sacha Gnjatic

## Abstract

Multiplex immunostaining analysis remains fragmented, underperforming, and labor-intensive despite tissue proteomic methodologies achieving ever-increasing marker complexity. Here we propose an open-source, semi-supervised automated pipeline that streamlines start-to-finish, single-cell resolution analysis of whole-slide tissue, named Multiplex-imaging Analysis, Registration, Quantification, and Overlaying (MARQO). We compared and validated MARQO using Multiplex Immunohistochemical Consecutive Staining on a Single Slide (MICSSS) using human tumor and adjacent normal tissue samples. Performance was compared with manually-curated pathologist determinations and quantification of multiple markers. We also optimized MARQO to analyze diverse tissue sizes (whole tissue, biopsy, tissue microarray) and staining approaches (singleplex immunohistochemistry, 20-color multiplex immunofluorescence) to determine marker co-expression patterns in multiple human solid cancer types. Lastly, we validated CD8 T cell enrichment in hepatocellular carcinoma responders to neoadjuvant cemiplimab in a phase II clinical trial, further demonstrating MARQO’s ability to provide spatially-resolved *in situ* mechanisms by providing multiplex whole-slide single-cell resolution data.

## INTRODUCTION

Multiplex immunohistochemistry (mIHC) and immunofluorescence (mIF) are essential imaging tools for determining single-cell resolution protein co-expression levels while maintaining spatial tissue integrity. Recent technologies such as CO-detection by inDEXing (CODEX)^1^, Cyclic immunofluorescence (CyCIF)^2^, multiplexed immunofluorescence^3^, Multiplexed Ion Beam Imaging (MIBI)^4^, and Multiplexed Immunohistochemical Consecutive Staining on Single Slide (MICSSS)^5^ have drastically increased the number of targeted proteins stained on one slide, creating many opportunities to better understand tissue organization and cell infiltration. In fact, characterizing and spatially understanding immune cell recruitment and organization in cancer lesions during treatment with immune checkpoint blockade (ICB) treatment has helped elucidate mechanisms of response or resistance and clarify interactions among cells, emphasizing the feasibility and advantage of multiplex imaging technologies^6^. While the experimental workflow of imaging technologies has been increasingly streamlined, the methodology for analyzing and producing whole-slide, quantitative multiplex data remains discrepant and computationally intensive, if at all possible, with available third-party analysis tools^7,8^. Although artificial intelligence is being investigated as a solution, proprietary algorithms make it difficult to accept outputs without confirmation from pathology review. Therefore, a pipeline that reliably streamlines and integrates whole-slide analysis for multiplex and singleplex IHC and IF using an automated but semi-supervised approach is needed.

Here we report the Multiplexed imaging Analysis, Registration, Quantification, and Overlaying (MARQO) pipeline and its validation by pathologists. It employs multiple interchangeable modules including dynamic deconvolution of channels, co-registration, segmentation of cell nuclei, and clustering of unique cell populations, all performed locally or in the cloud. Each step has adjustable parameters that have been optimized and can be adjusted for diverse tissue types and staining protocols (MICSSS^5^, COMET IF^9^, Orion^10^, Vectra^11^). Other recent multiplex analysis tools focus on specific aspects of the quantification methodology, such as co-registration or nuclear segmentation^12^, requiring the user to toggle across applications and learn new software. This fractionation of software is a hurdle for harmonized quantification across platforms, an essential requirement of large networks for correlative science such as CIMAC-CIDC^13^. MARQO (1) utilizes the iterative qualities of multiplex data to help denoise and refine repetitive data, utilizing the multiplex aspect of the assay to its advantage. Moreover, while existing tools are mostly released as GitHub repositories, only operational through the Command Prompt interface, MARQO (2) is easily operational to both computational and non-computational users via a Command Prompt interface as well as a graphical user interface (GUI). Lastly, we have employed a “semi-supervised” approach to characterize cell populations, because fully automated decisions about positive signals are not yet acceptable clinically without pathologist confirmation. The approach of unsupervised clustering followed by a supervised binarization of each cluster by the user permits MARQO to (3) accurately quantify and cluster cell populations across multiple platforms, tissue types, and markers of interest without the need to train the machine-learning (ML) algorithms while still permitting closely monitored quality control by the user.

We applied MARQO to whole-slide MICSSS data, a technology developed and broadly used by our laboratory^14–16^, but difficult to analyze with existing tissue analysis tools (HALO^17^, Visiopharm^18^, QuPath^19^), to study spatial immune responses to ICB in cancer. While neoadjuvant ICB targeting the PD-1/PD-L1 axis has revolutionized treatment approaches for various cancer types^20^, the response to immunotherapy can vary in success^21^ and many patients remain left out. CD8 T cell enrichment has been associated with the response to ICB^22–24^. Leveraging a trial of neoadjuvant cemiplimab (anti-PD-1 antibody, ClinicalTrials.gov, <u>NCT03916627</u>, Cohort B) in patients with hepatocellular carcinoma (HCC)^25^, we recently corroborated these findings in responders to ICB. However, to fully understand the mechanisms driving the response to treatment, we assessed multiple cell populations using MICSSS and used MARQO to resolve the multiparametric cellular and spatial data in responders to ICB relative to non-responders. With MARQO, we detected CD8 T cell enrichment in specific tissue areas defined by a pathologist as tumor, fibrotic, and necrotic. In addition, we analyzed immune cell organization and proximities in tumors pre-and post-treatment. As a result of this novel method of multiplex imaging and analysis, we are better able to elucidate the quantitative mechanisms that underlie the responses and resistance to neoadjuvant anti-PD-1 treatment.

## RESULTS

### MARQO enables analysis of diverse imaging technologies

MARQO was tested and validated for its ability to analyze MICSSS, mIF (COMET IF), and singleplex IHC (multiple chromogens), using the workflow described in Figure 1. Following each respective protocol for producing and scanning all stained tissue slides, MARQO can analyze these datasets locally or via the Cloud. To begin the analysis, the user operates the Command Prompt or downloadable GUI to quality control each sample by specifying input data types, including the type of assay and the name of each stain (Supplementary Video 1). This is followed by a tissue masking and, if applicable, a preliminary registration step, to ensure that the tissue is being correctly captured. The next steps of the pipeline run automatically and depend on the type of data being analyzed. For example, while IHC technologies require a color deconvolution to separate the chromogen from the nuclear counterstain (usually hematoxylin), multiplex IF images contain a single stain per channel in a multidimensional array, which usually requires a denoising step. The pipeline divides the tissue into evenly sized tiles, allowing parallel computational analysis and reducing computation time. On a per-tile basis, the registration, segmentation, and quantification modules are enacted, again dependent on the technology being analyzed. While some mIF technologies require the registration step^2^, others are already aligned and can forgo this step^9^. This step is followed by a clustering module, in which each stain is treated as an independent input, and segmented cells are compartmented into multiple clusters. The user is subsequently prompted to perform another quality control to review the performance of the analysis and binarize the positivity of each cluster per marker.

**Figure 1:**
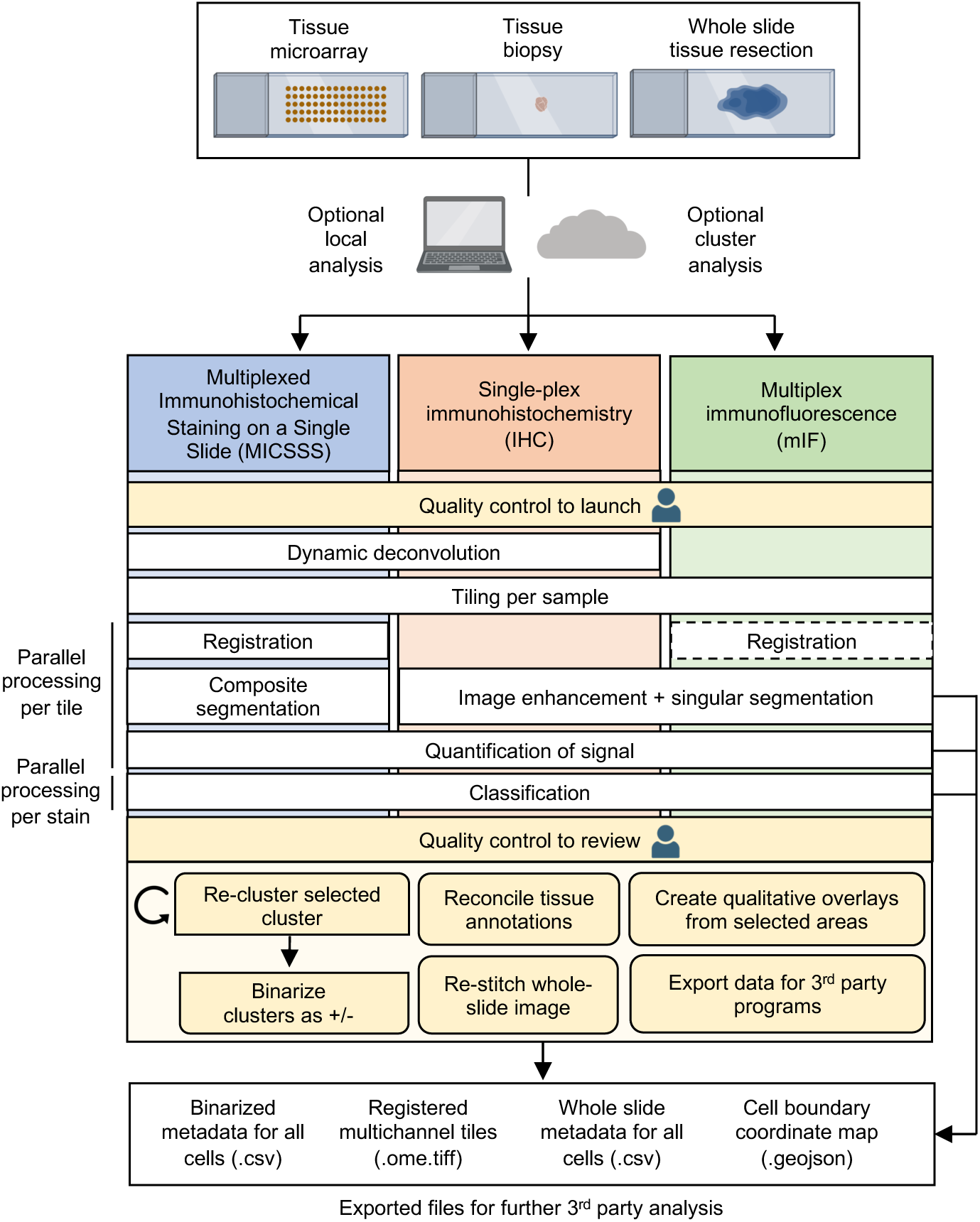
MARQO pipeline overview. Variable FFPE tissue sizes from small TMA cores to whole-slide tissue resections can be analyzed utilizing a tiling architecture, which can be performed locally or externally via cluster or cloud. The pipeline is compatible for slides stained with MICSSS, singleplex IHC, or mIF. After the user provides a brief quality control when importing data, the pipeline independently processes each staining technology differently. MICSSS and IHC undergo dynamic deconvolution to extract nuclear and chromogen staining. All technologies are converted into numerous smaller, overlapping tiles for faster, parallel processing. The multiplex technologies undergo a series of registrations. All technologies are segmented for cell nuclei, where MICSSS uses a composite methodology from its iterative nuclear staining. All technologies quantify the chromogenic positivity staining, nuclear staining, and morphological information. These metadata are used for cluster-based classification, after which the user quality controls to assess which markers the clusters truly stain positively or negatively for. The user can use MARQO downstream analysis tools to further review data. These modules can reconcile tissue compartment annotations from QuPath, re-create large chromogenic or whole-channel image files that combine multiple individual tiles, create figure-ready qualitative overlays to visualize cell populations, and export data for third-party programs. At multiple checkpoints in the pipeline, intermediary files that are also easily integrated into third-party programs are exported.

Numerous benchmarking data outputs are generated, not only to provide the user with closely monitored progress reports for each step for inspection, but also so that the user can conveniently toggle across different analysis platforms for continued analyses (Supplementary Table 1).

### Iterative staining of cells is harnessed to improve the composite segmentation mask

For MICSSS and some mIF, repetitive nuclear staining can be harnessed to further refine and denoise imaging data. While some mIF technologies use DAPI for nuclear counterstaining, MICSSS counterstains nuclei with hematoxylin per staining cycle iteration. For this reason, we stress the denoising benefits of multiple nuclear or cytoplasmic stains, while using MICSSS in our validation. MARQO performs a novel registration technique for MICSSS in which each tile per iterative stain is matched to the respective tile of the first stain that encompasses roughly the same tissue area. This process results in parallel batches of tiles that are aligned by their deconvoluted nuclear stains, which is assumed to remain consistent across each batch since there is no chromogen or other-colored artifact in this channel (Figure S1). MARQO then performs a supervised nuclear segmentation iteratively across stains per batch of tiles (Figure 2A). Here, the pipeline harnesses the power of multiplex nuclear staining to further refine its segmentation mask. If the centroid of a nuclear object is consistent within a tunable margin across the series of stains (defaulted at 60% of stains), it is kept in the final composite segmentation mask. Otherwise, we assume these objects are artifacts improperly classified as nuclei within an individual staining or represent cells that were eliminated during iterative staining due to tissue damage. Similarly, based on staining and available metadata, the pipeline further refines the mask by eliminating hypothesized red blood cells (RBCs) (Figure S2A).

**Figure 2:**
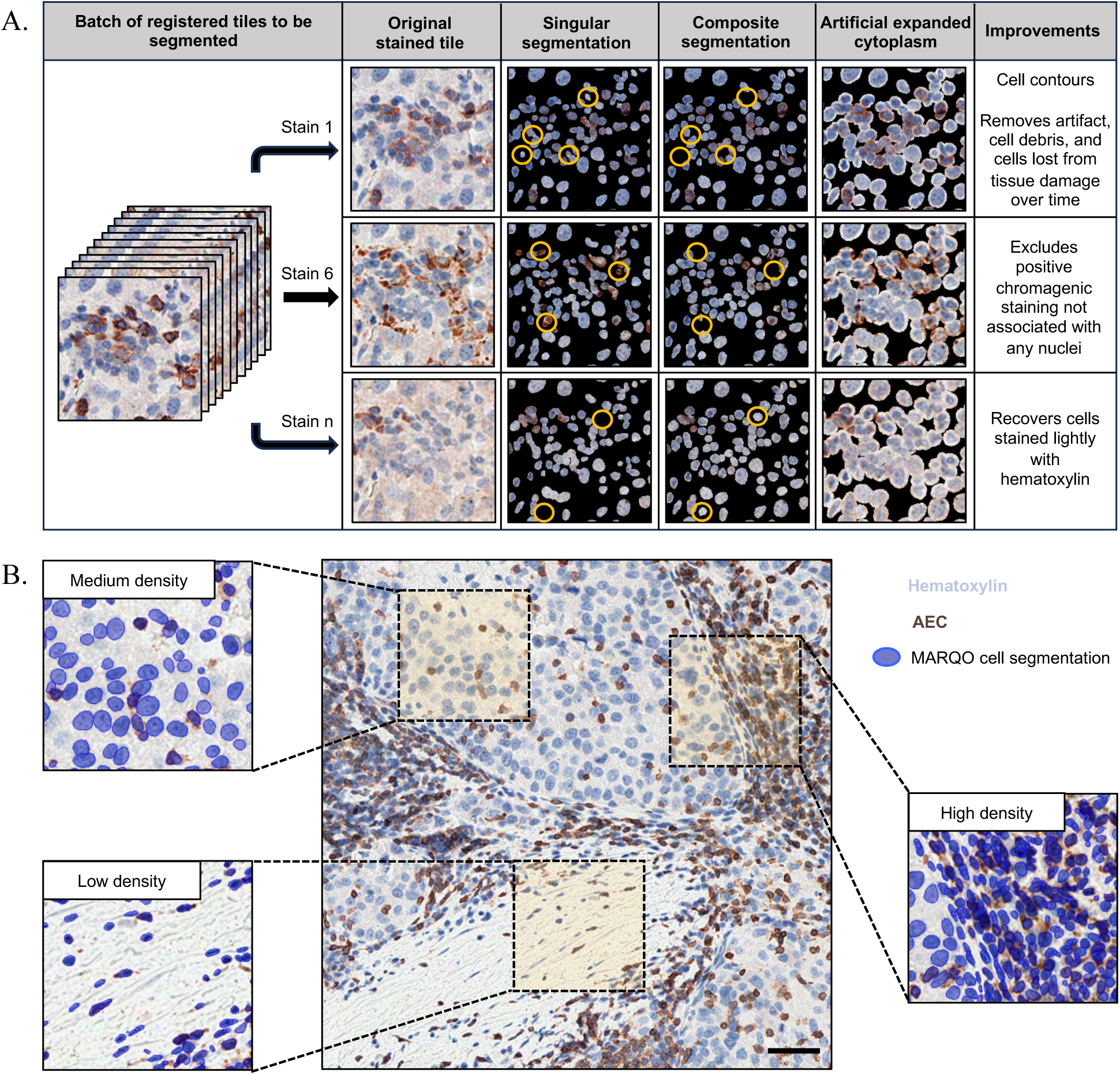
Detailed singular and composite segmentation. **A)** An example batch of tiles that spans 1-*n* stains from MICSSS undergoes novel composite segmentation. The first column shows the original staining prior to segmentation, then the nuclear segmentation mask if each marker were independently segmented, then the nuclear segmentation from reconciling multiple nuclear stains, followed by the composite segmentation with nuclear boundaries extended by 3 pixels to simulate cytoplasm. The last column provides examples of the improvements that composite segmentation has over the singular segmentation. **B)** An example tile of MICSSS depicts MARQO composite segmentation across diverse cell densities, showing with zoom-ins how MARQO similarly segments in low-, medium-, and high-density areas. Scale bar is 50µm. Comparisons with QuPath segmentation are provided in Supplementary Fig. S2.

To assess and validate the performance of MARQO’s composite segmentation, we used samples stained with an onco-immune targeted MICSSS panel. We analyzed a cohort of patients with HCC who received neoadjuvant cemiplimab treatment followed by surgical resection; for these patients, we analyzed pre-treatment biopsies and HCC resections. Using 10 500×500µm region-of-interests (ROIs) from these tissues, we compared the composite segmentation performance of MARQO to that of semi-manual segmentation by a pathologist using a conventional third-party analysis tool, QuPath^19^. We found that MARQO’s segmentation performed well across heterogeneous densities, which is usually problematic because cell segmentation is set up for a global and homogeneous density (Figure 2B). Overall, MARQO segmented ±10% of total cells determined semi-manually across the ROIs and achieved an average Sørensen–Dice coefficient (DICE score) of 0.83 using an Intersection over Union (IoU) threshold of 0.5, implying that if a semi-manually segmented cell and a MARQO-segmented cell overlapped by a conventionally-accepted amount of 50%, an average of 83% of cells in all evaluated ROIs were segmented identically in QuPath and MARQO (Figure S2B-E). In accordance with recent guidelines for including tissue variability^26^ and pathologist supervision^27^ for improving automated pathology algorithms, these data support the claim that MARQO segments similarly, if not better, to the semi-manual segmentation performed in QuPath.

### Rapid clustering and re-clustering of cell populations improve quality control

Once a batch of tiles representing an entire tissue section is analyzed, a features table of segmented cells is appended with metadata associated with all segmented cells within that region, which provides the framework for the classification module. While many current classification approaches leverage supervised machine learning, which requires previously-trained datasets and potentially introduces bias^28^, we envisioned a model that could be applied across multiple technologies, tissue types, and markers without additional training. We also wanted to permit close interpretation of the results, since learning models are not mature enough to be used entirely without human supervision in digital pathology^27^. MARQO combines unsupervised clustering for all cells and supervised classification per marker for each cluster by the user, herein called a semi-supervised approach. MARQO performs a faster clustering approach for larger samples (minibatch K-means), as well as a more intensive, but lengthier clustering approach for smaller samples (Gaussian mixture model), if needed. By default, and for the analyses in this paper, MARQO uses the minibatch k-means algorithm. When the automated clustering finishes, the user operates a GUI to assess which of the produced clusters of cells are positive or negative per marker (Figure 3A, supplementary video 2). If the user deems a certain cluster still mixed with positive and negative cells, the user may “re-classify” a cluster to output a new round of sub-tiers within the original tier, permitting the user to closely fine-tune classification outcomes.

**Figure 3:**
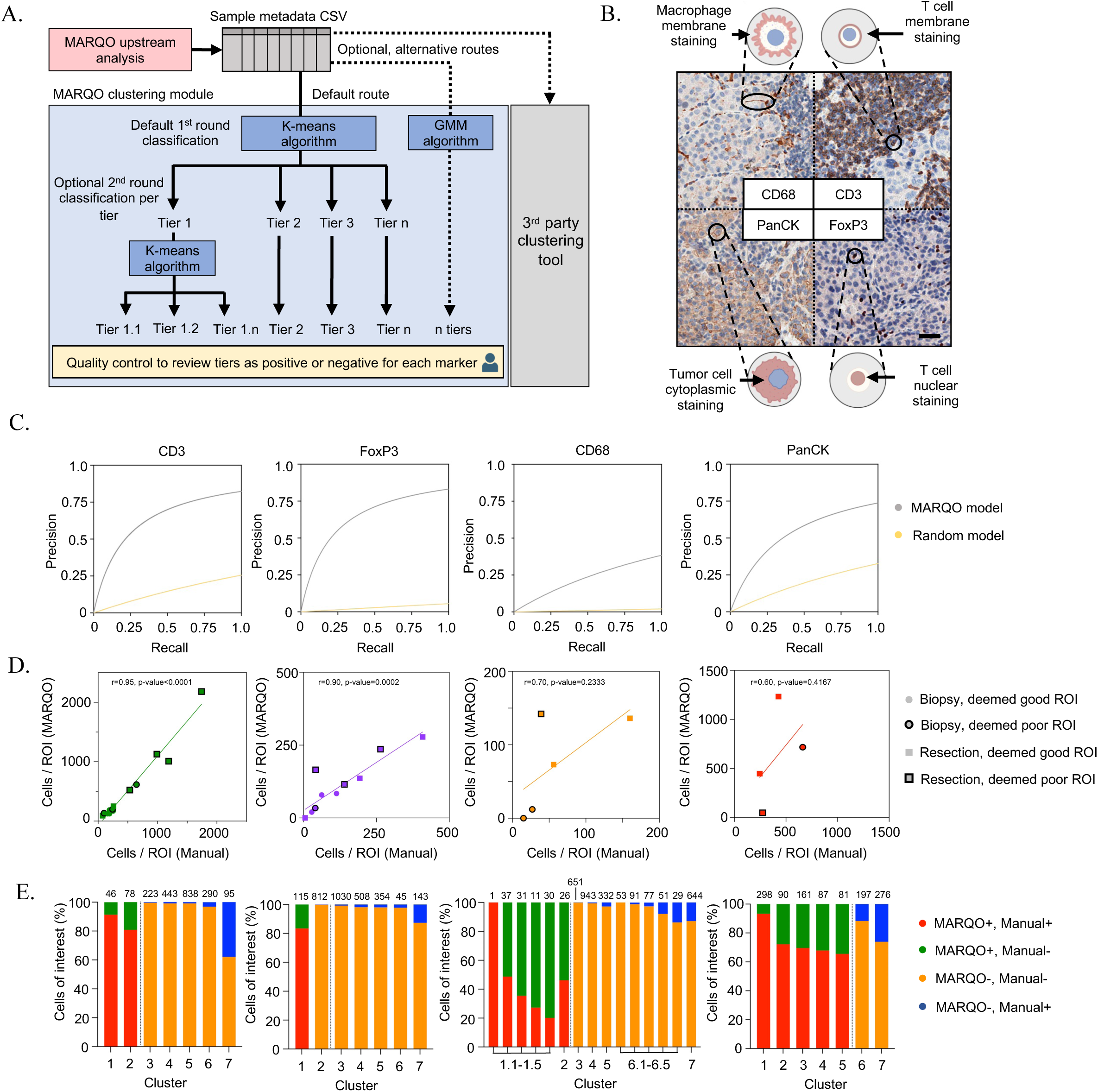
MARQO classification with validation. **A)** Following upstream analysis and production of a metadata CSV per sample, MARQO can classify cells via the default K-means algorithm or alternative routes such as Gaussian Mixed Model (GMM) clustering and third-party tools. Following K-means or GMM routes, MARQO categorizes cells into a predefined number of clusters, which the user can inspect in the MARQO application. If needed, the user can split a cluster into sub-clusters, which rapidly performs K-means classification for the chosen cluster. The user can inspect all clusters and sub-clusters to assess whether they are positive or negative for each marker. **B)** To validate our classification approach, we chose four diverse markers: nuclear marker FoxP3, circular shape membrane T cell marker CD3, asymmetrical shape membrane marker CD68 (macrophages), and tumor cytoplasmic marker PanCK. Scale bar = 50 µm. **C)** Precision versus recall graphs for CD3, FoxP3, CD68, and PanCK comparing MARQO classification performance to that of pathologist manual positivity assessment (predictive model); a random model is also graphed. **D)** Scatterplots with regression analysis lines comparing total number of cells manually deemed positive by the pathologist versus by MARQO across 34 total ROIs and the four chosen markers, with each point signifying biopsy or resection tissue and good- or poor-quality staining, deemed by the pathologist. Spearman correlation r values with affiliated p-values are shown per marker. **E)** Stacked bar graphs depicting a representative ROI for each of the four chosen markers. Whereas the user quality controls classified clusters per sample, here we show how that breakdown appears per the chosen ROI that the pathologist also manually annotated. The colors within each cluster define the proportion of cells that were deemed positive or negative by the user on MARQO or by the pathologist. Numbers above the bars denote the total number of cells within that cluster. CD68 is an example of when the user re-classified the data into sub-clusters. The corresponding ROIs are provided in Supplementary Fig. S3.

To validate the performance of our classification module, we leveraged our cohort of resected HCC samples and identified four markers that have unique challenges when quantifying with currently available platforms: nuclear marker FoxP3 (regulatory T cells, Treg), circular shape membrane marker CD3 (T cells), asymmetrical shape membrane marker CD68 (macrophages), and cytoplasmic marker PanCK (tumor cells, Figure 3B). We compared MARQO positive classification results after user quality control to the manually counted positive cells by a pathologist. We used 34 ROIs from different samples used in the segmentation validation, which were also deemed as good- or poor-quality areas by the pathologist based on the degree of tissue damage and staining artifacts. Using the manually counted cells as the predictive model, MARQO achieved 83% specificity and a Spearman correlation coefficient r=0.95 (p<0.0001) for CD3 and 90% specificity and r=0.90 (p=0.0002) for FoxP3 (Figure 3C-E, S3). It also achieved 97% specificity and r=0.70 (p=0.2333) for CD68 and 85% specificity and r=0.60 (p=0.4167) for PanCK, markers that are conventionally difficult to quantify due to their amoeboid-like shapes and heterogeneous staining. We also visualized the performance of MARQO classification for additional markers, including PD-1, CD8, Ki-67, αSMA, CD20, and MZB1 (Figure S4). These data support the claim that MARQO’s fast semi-supervised classification performs similarly to the lengthy, manual annotations performed by a pathologist, which grants the user time and ability for continued downstream analysis.

### Diverse tissue sizes and types are processed using MARQO

MARQO can theoretically process any-sized tissue due to its tiling architecture. However, during multiplex staining, different-sized tissue sections possess unique biological, technical and computational challenges: smaller tissues have increased relative edge effect staining^29^ and may incidentally contain rare, niche cell populations, which are less likely to be isolated during clustering; meanwhile, larger tissues are more prone to tissue damage^30^, frequently suffer from tissue folding, and require vast computational resources. To assess whether MARQO performs similarly across different-sized tissues, we analyzed tissue microarray (TMA) cores, core needle biopsies, and samples from surgical resections that were processed with MICSSS (Figure 4A). Moreover, we assessed and validated classification consistency across diverse solid tumor types, since diverse tissues can contain unique morphologies and marker expression patterns^31^ (Figure 4B). Following MARQO analysis and user review quality control, we determined marker densities for CD3 and PanCK. The cohort included whole slide resected tissue of non-small cell lung cancer (NSCLC) and TMA cores from the following tissue types: NSCLC, head and neck squamous cell carcinoma (HNSCC), colorectal cancer (CRC), breast cancer (BC), epithelial ovarian cancer (EOC), pancreatic ductal adenocarcinoma (PDAC), glioblastoma (GBM), renal cell carcinoma (RCC), and melanoma (MEL). We compared densities generated by MARQO or by a pathologist through a standardized methodology on QuPath^19^. We determined that CD3 had an overall Spearman r=0.98 (p<0.0001) and PanCK an overall r=0.90 (p<0.0001) across all tested tissues (Figure 4C). These findings suggest that MARQO performs similarly to the manual quantification by pathologists for diverse tissue sizes and types.

**Figure 4:**
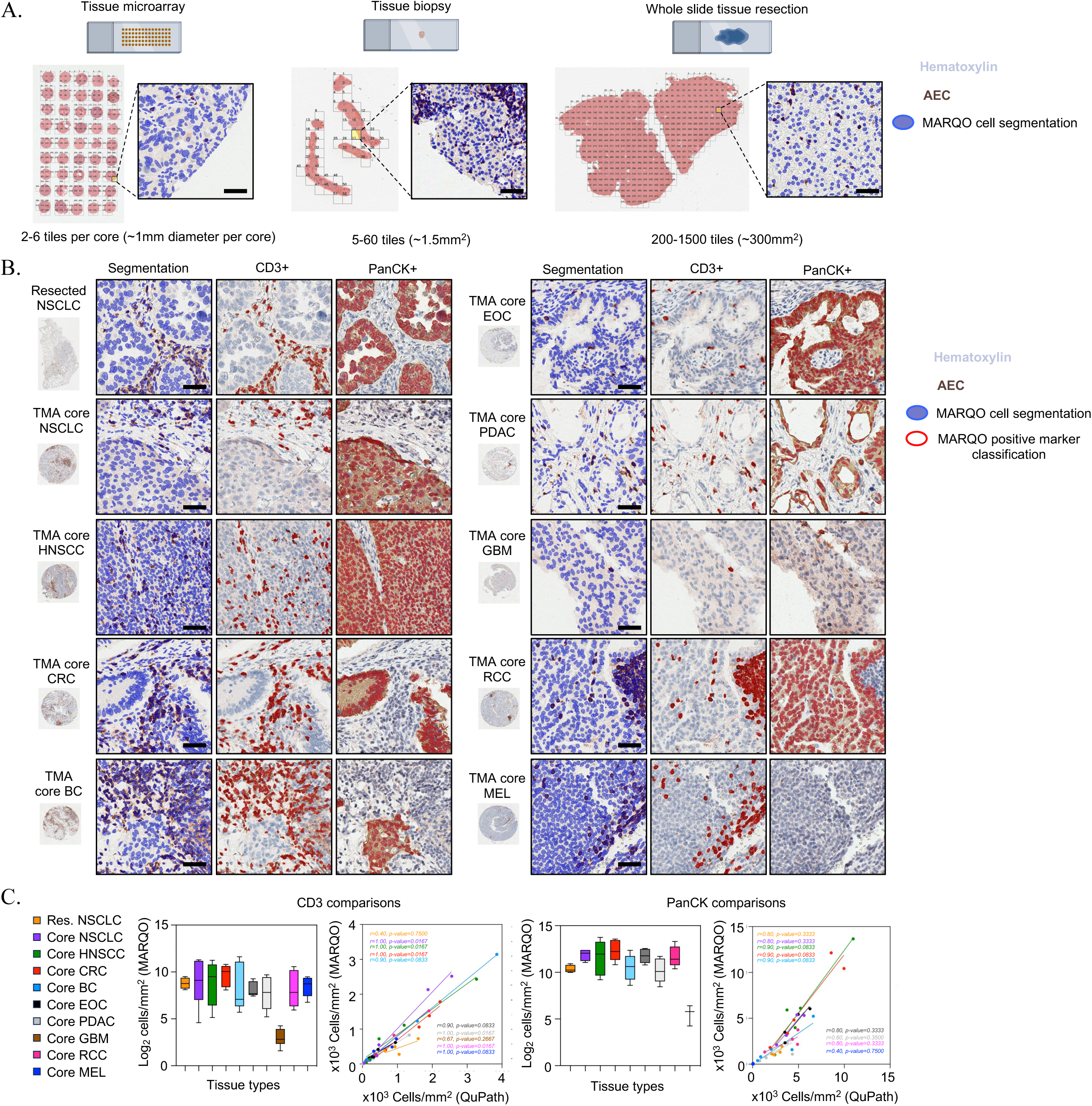
Validation across diverse tissue sizes and types. **A)** We tested small TMA cores, tissue biopsies, and whole-slide tissue resections on MARQO, with these tissues tiled and segmented here. Scale bars = 50 µm. **B)** MARQO’s segmentation and classification performance after quality control by the user for markers CD3 and PanCK are shown for the following tissue sizes and types: whole slide resected tissue non-small cell lung cancer (NSCLC) and TMA cores for NSCLC, head and neck squamous cell carcinoma (HNSCC), colorectal cancer (CRC), breast cancer (BC), epithelial ovarian cancer (EOC), pancreatic ductal adenocarcinoma (PDAC), glioblastoma (GBM), renal cell carcinoma (RCC), and melanoma (MEL). Scale bars = 50 µm. **C)** Box plots depicting MARQO densities and scatterplots with regression analysis lines comparing MARQO densities to QuPath-derived densities for the corresponding tissue sizes and types are shown here for markers CD3 and PanCK. Spearman correlation r values with affiliated p-values are shown per tissue type per regression.

### MARQO integrates workflow for multiple staining technologies

Next, we assessed MARQO’s performance to analyze diverse types of assays. While MARQO has been optimized for quantifying cell subset density based on consecutive stains and multiple markers, we assessed its ability to analyze singleplex IHC, an assay often used for routine staining of markers of interest. We compared marker densities for CD3-stained cells with another commonly used revelation reagent, chromogen diaminobenzidine (DAB), and used the same reagent used for MICSSS, chromogen aminoethyl carbazole (AEC), to look at PD-L1, a marker used in the clinic for diagnostic and prognostic purposes^32^. While we acknowledge that this validation is for research purposes only, we demonstrate that with further clinical validation, MARQO may be employed in a clinical setting. We determined densities acquired by a pathologist with QuPath on an independent cohort of 5 samples. For CD3, we determined a Spearman r=0.60 (p=0.3500) and for PD-L1 we determined r=1.00 (p=0.0167) for positively classified cells between MARQO and QuPath (Figure 5A). Similarly, to confirm the accuracy of MARQO for segmenting and characterizing cell populations for non-IHC multiplex technologies, we analyzed 4 whole-slide NSCLC samples via COMET IF, a technique utilizing mIF. We compared cell densities of FoxP3, CD3, CD68, and PanCK between MARQO and QuPath, obtaining an overall r=0.92 (p<0.0001) for all the markers (Figure 5B). In addition to testing and validating singleplex IHC and COMET IF, we also tested other platforms on MARQO, including Orion^10^, CODEX^1^, and CyIF^2^ (Supplementary Table 1). Like our previous results, these data suggest that MARQO retains high accuracy when analyzing diverse tissue and assay inputs.

**Figure 5:**
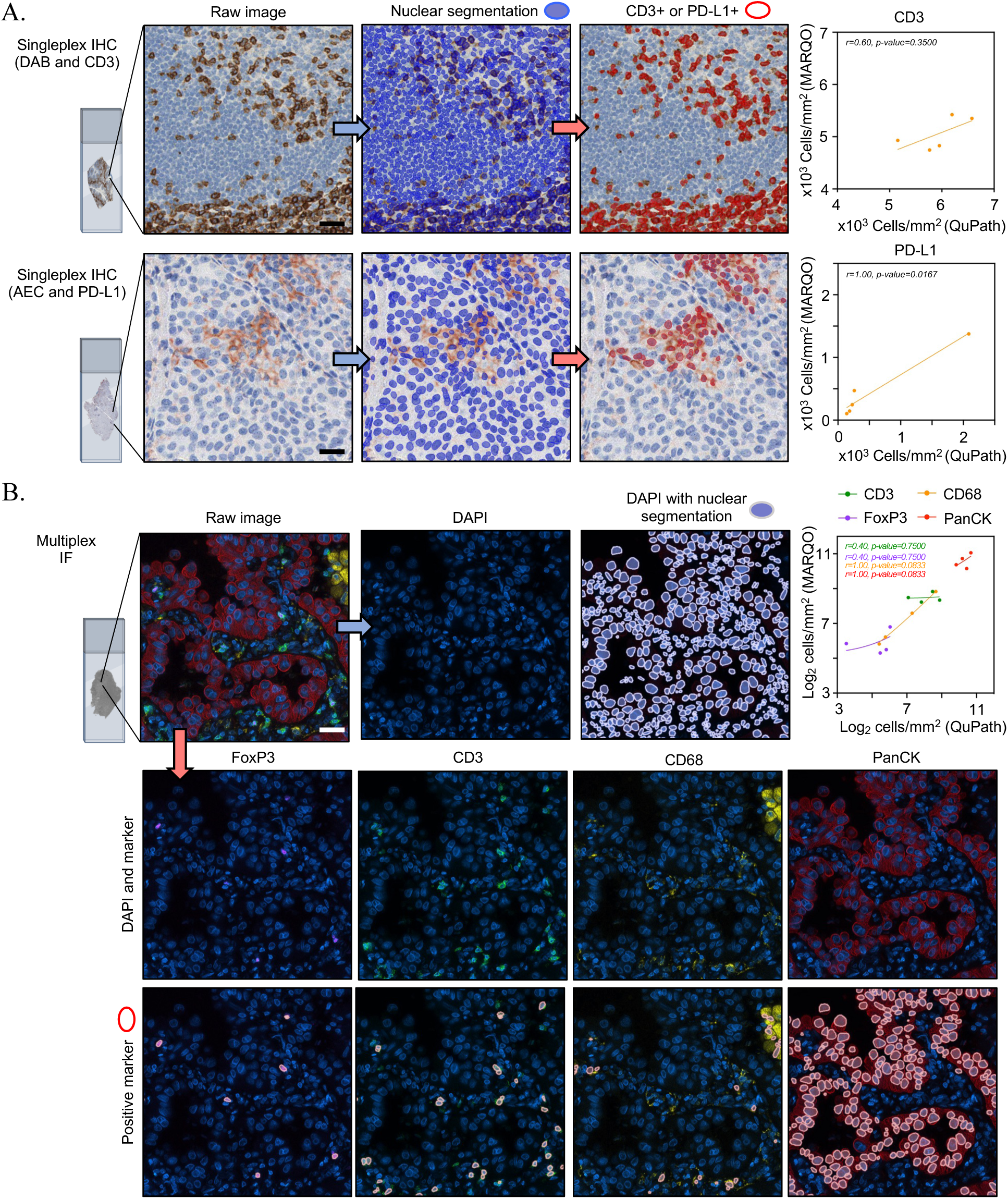
Integrative platform for other staining technologies. **A)** Singleplex IHC examples segmented and classified after quality control by the user for marker CD3 with chromogen Diaminobenzidine (DAB) (top) and marker PD-L1 with chromagen aminoethyl carbazole (AEC) (bottom). Scatterplots with regression analysis lines compare MARQO densities to QuPath-derived densities for CD3 (top) and PD-L1 (bottom). Spearman correlation r values with affiliated p-values are shown per marker. Scale bars = 30 µm. **B)** mIF using COMET IF technology, to assess MARQO’s ability to segment and classify another multiplex technology after user quality control. We depict a representative tile with cellular nuclear stain DAPI and the four previous overlapping markers of interest: FoxP3, CD3, CD68, and PanCK. Stemming from this tile (right) is the single DAPI stain, adjacent to the MARQO segmentation mask overlaid on the DAPI image. Also stemming from the tile (bottom) is each of the marker channels of interest above that marker channel with MARQO’s classification channel after user quality control overlaid. Top right panel is a scatterplot with linear regression analysis lines to compare MARQO densities to QuPath-derived densities for all four markers. Spearman correlation r values with affiliated p-values are shown per marker. Scale bar = 30 µm.

### MARQO identifies CD8 T cell enrichment in responders to neoadjuvant cemiplimab treatment

Multiple downstream analyses can be performed in the MARQO GUI from the final summary table, which can facilitate further quantifications and analyses (Supplemental Video 2). We leveraged a clinical trial cohort of 18 patients with early-stage HCC treated with neoadjuvant cemiplimab and used MARQO to better understand the immune cell infiltration in responders (Resp, n=6) versus non-responders (NR, n=12). We used MICSSS and MARQO to analyze pre-treatment HCC biopsies (baseline) and HCC surgical resections after two doses of cemiplimab (post-treatment) and observed that not only did the response to cemiplimab modify the global immune infiltration compared to NR, but the immune landscape was already different at baseline between these patients (Figure 6A). A pathologist demarcated tumor, non-tumor adjacent liver (adjacent), fibrotic, and necrotic regions in each tissue section using QuPath to understand the specific cell distribution among the samples (Figure 6B-C). We found that, unsurprisingly, NRs were mostly represented by tumor tissue, whereas this compartment completely disappeared in Resp post-treatment. Specifically, we were interested in CD8 T cells, since we previously characterized their clinical relevance and spatial interactions in response to cemiplimab treatment^33^. Using our GUI, we reconciled the QuPath annotations with cell classification, producing a table that identified the location of each cell within tumor, adjacent, fibrosis, and necrosis. We detected enrichment of CD8^+^CD3^+^ T cells in fibrotic and necrotic areas of post-treatment Resp compared to baseline (p=0.3398 and p=0.0267, respectively) (Figure 6D). Interestingly, we found that CD8 T cells were already enriched in Resp at baseline compared to NR (p=0.008). To further explore this phenomenon, we used a neighborhood analysis approach to determine the shortest distance between cell types of interest per cell and how the distance varied with treatment in the different annotated areas. We found that CD8 T cells tend be very close to each other in Resp compared to NR in fibrotic and necrotic regions both at baseline (p=0.0012 and p=0.0187, respectively) and post-treatment fibrosis (p=0.0162), suggesting immune aggregations (Figure 6E). In comparison, CD8 T cells were generally found within 50 µm of B cells and Tregs, suggesting a potential interaction. In summary, these results demonstrate the quantitative metrics available using MARQO and provide the first example of cellular and spatial reconciliation of cancer lesions using MICSSS.

**Figure 6:**
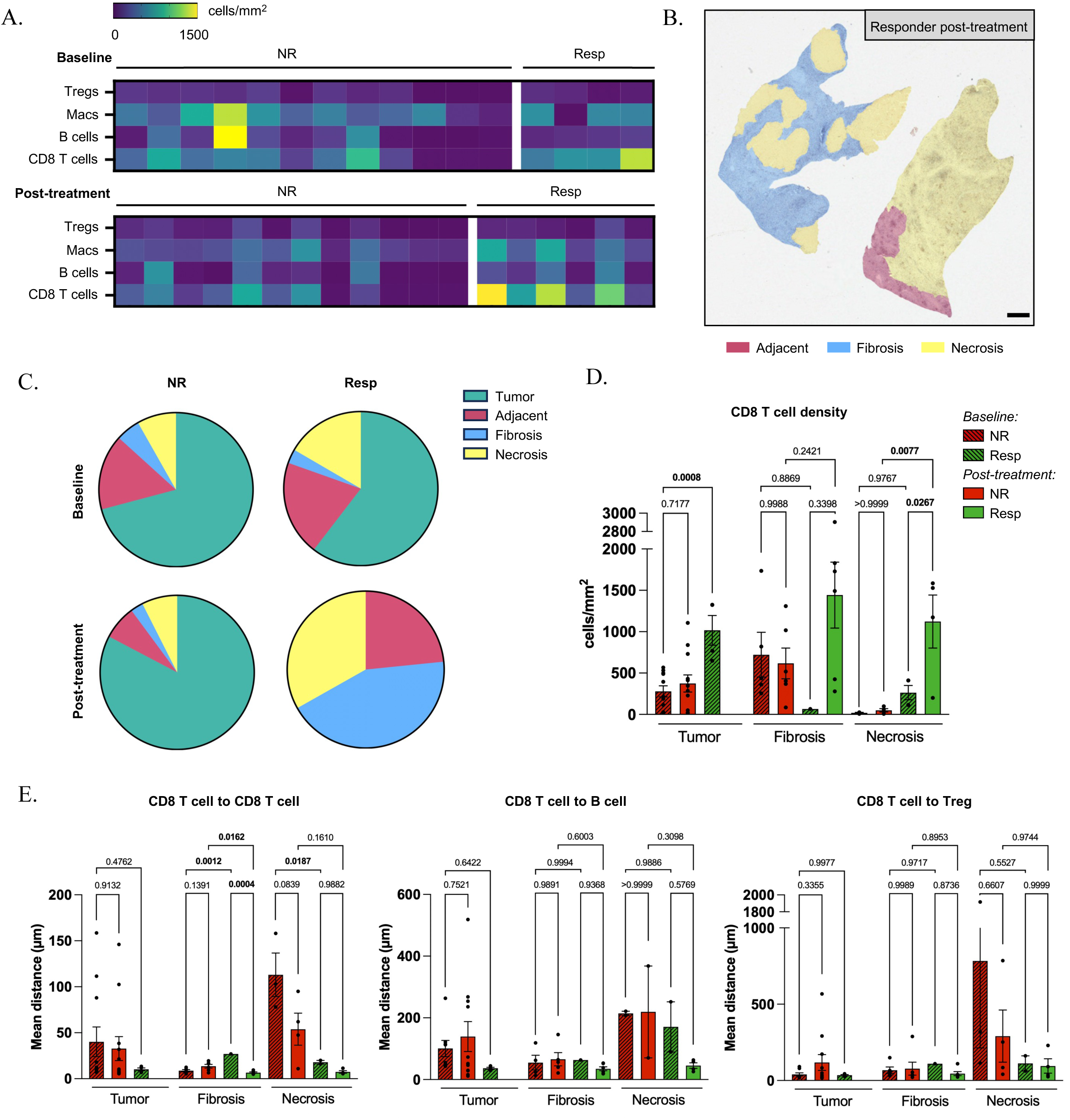
MARQO identifies CD8 T cell enrichment in responders to anti-PD-1 treatment. **A)** After classification and review, we identified Tregs (CD3+FoxP3+CD8-), Macs (CD68+), B cells (CD20+), and CD8 T cells (CD3+CD8+) and evaluated their densities across whole-slide tissues for responders (Resp) and non-responders (NR) to cemiplimab treatment at baseline and post-treatment. **B)** Representative image showing the overlay of the tissue with the annotation made by the pathologist using QuPath to demarcate tissue region boundaries, including tumor, adjacent, fibrosis, and necrosis, for all tissues used. Scale bar = 500 µm. **C)** Pie charts depicting the tissue compartments annotated by a pathologist for all samples at baseline and post-treatment for Resp (n=4 and 6 respectively) and NR (n=12 for both). **D)** CD8 T cell densities plotted at baseline and post-treatment for NR and Resp groups within tumor, fibrosis, and necrosis compartments. **E)** Bar plots depicting the means of the shortest distances from CD8 T cells to themselves (left), to B cells (middle), and to Tregs (right) at baseline and post-treatment for NR and Resp within tumor, fibrosis, and necrosis compartments.

## DISCUSSION

Here we introduced and validated the MARQO pipeline to quantitatively analyze whole-slide singleplex IHC, mIHC, and mIF, enabling quantitative interpretation of cellular and spatial organization in cancer tissue lesions. Through its diverse modules and adjustable parameters, MARQO can be tailored to analyze data from multiple platforms, tissue types, and markers of interest, outputting unprecedented co-expression whole-slide data using a single platform. MARQO and its intermediate diagnostic files seamlessly integrate with third-party downstream analysis tools. Moreover, MARQO provides a GUI-based module that formats the summary outputs to the software of interest. Specifically, MARQO provides the first throughput whole-slide analysis tool for MICSSS, which was, until now, suboptimal or labor-intensive using standard tools such as QuPath^19^, Halo^17^, or Visiopharm^18^ due to the iterative nature of the staining protocol and need for precise image alignment. Rather than using an artificial intelligence approach that would require extensive training for diverse tissue types without a means to confirm specificity, we deployed a “semi-supervised” approach to cluster and classify cell populations using dynamic thresholds and user-controlled selection of positive versus negative signals. This approach can be performed per sample or per group of selected samples, but further validation and testing must be performed to validate the accuracy of additional markers and tissues, as well as multiple samples classified at once. While validation has been performed for the samples presented here, we remain confident that MARQO can deliver robust data for diverse preclinical and clinical tissues across a range of pathologies.

An inherent limitation of whole-slide IHC and IF analysis is the size of the required computational resources. While we and other analysis tools^8,12^ have utilized a tiling algorithm to parallellize processing and reduce computation time, multiple central processing units (CPUs) are needed, which are not always available with local resources. Considering this, biopsy and TMA analysis are feasible locally, but we strongly recommend Cloud resources for whole-slide multiplex imaging analysis. Another limitation includes the quantification of individual staining as binary outputs, and cell identification and segmentation based on nuclear detection with artificial cytoplasmic extension. Intensity scoring could be considered for appropriate staining protocols in future versions, based on expected tier differences during selection of similarly stained cell clusters, and alternate segmentation protocols could be integrated based on membrane markers when available. These areas of improvement, as well as additional tools to further analyze multiplex imaging data, are our current targets for future deployments and updates to the available pipeline, which can easily be appended as additional modules in our application.

We envision that MARQO will drive quantitative discoveries in imaging assays. As the first tool to analyze whole-slide MICSSS, we used MARQO to quantify enrichment and localization of CD8 T cells in HCC responders to neoadjuvant cemiplimab. These findings elucidate previous hypotheses about the interactions of CD8 T cells and will help us better understand mechanisms of response or resistance to immunotherapy^33^. Spatial biology is emerging as a pivotal field for biomarker discovery. By understanding tissue organization, immune cell infiltration, and antigen presentation heterogeneity in a harmonized manner, researchers can interpret data across multiple studies and platforms, ultimately developing actionable predictive tests^34–37^. With MARQO, we hope pathologists and researchers will have a necessary and novel tool to quantitatively analyze whole-slide imaging data, augmenting future discoveries and bolstering the field of cancer immunology.

## MATERIAL AND METHODS

### MARQO modules

#### Operating the MARQO pipeline

The MARQO pipeline can be deployed through different container technologies, such as Docker and Singularity. The container is built on the top of Tensorflow’s pre-built docker image, where a python virtual environment contains all the libraries it requires. The container can be executed virtually in any kind of environment: single or multiple cores, single or multiple computers, servers, high performance computing clusters, or personal computers. The user has the option to interact with a web-based GUI or work through a command line interface via the Command Prompt, where they will be able to use a semi-automatic approach to fine-tune parameters for both the launching and reviewing features (Supplementary videos 1 and 2).

In the launching tab, the user selects the imaging technology to analyze and then executes tissue masking, preliminary registration, tiling (image decomposition into smaller images) and color deconvolution. While these steps are automated, the user may change their settings and rerun them for optimal quality control. For tissue masking, the user may draw a specific tissue mask to overwrite any automation, if deemed necessary. In the tiling step, the user provides input on how the algorithm should produce overlapping tiles, including how much area from the edges of the tissue should be excluded from analysis because of edge-case staining and folding and the minimum proportion of tissue each tile should contain to be analyzed. The channel deconvolution module is a feature from the HistomicsTK package^38^, which implements a revised method from Macenko *et al*.^39^. It performs a dynamic color deconvolution, extracting three channels based on a deconvolution matrix determined from the thumbnail-resolution whole-slide image for every stain. For MICSSS, this method extracts the chromogen stain, the nuclear hematoxylin counterstain, and a residual channel. A configuration file is created by saving the outputs for each of these steps so that the user may use the same quality control for future pipeline runs.

In the reviewing tab, the user may perform several downstream tasks in addition to classifying cell clusters, such as stitching together registered whole-slide images as RGB images or multi-channel arrays containing only positive staining. The user may also reconcile tissue compartment annotations to analyze cell infiltration in specific areas of interest, which can be obtained from an extracted and uploaded standardized “.geojson” containing the tissue compartment metadata. This will populate the final summary table by identifying what tissue area each cell resides in. The user can visualize cell populations against a backdrop of choice by toggling multiple positive and negative marker combinations via the live-updated GUI and obtain direct cell subset quantification and density for the whole tissue as well as each compartment annotated. Finally, the user may convert MARQO file outputs to third-party friendly formats for subsequent analysis and production of figure-ready qualitative overlays (Supplementary table 1).

#### MARQO whole-slide registration module

MARQO performs a series of registration steps for technologies requiring alignment to allow multiplex cell-resolution quantification. During the user’s initial quality control to launch a job, they performed a preliminary translational registration of the thumbnail low-resolution images, yielding relative alignment of tissues across iterative stains. Following the tiling module, batches of tiles will have 10% overlapping borders with other neighboring batches of tiles to permit registration and segmentation near tile borders without cropping cell boundaries. Per batch, the MARQO registration module deconvolutes all the tiles to extract the nuclear counterstains, since the counterstain remains consistent across staining iterations, unlike the tissue architecture or positive staining. The deconvolution matrix was dynamically determined prior in the launch application. MARQO then performs affine and elastic registrations to align all n^th^-stained tiles per batch to the 1^st^-stained tile, using the SimpleElastix^40^ registration package and parameters that were optimized using MICSSS images. The registered vector field is applied to the red, green, and blue channels for each tile. This process results in cell-resolution registration per batch of tiles.

#### MARQO segmentation module

For each batch of tiles registered, MARQO enacts its segmentation module to find cells within the tile so that it can later acquire each cell’s metadata in its quantification module. For technologies with iterative nuclear stains, such as MICSSS, the segmentation module iteratively performs semantic nuclear segmentation via STARDIST, a pre-trained algorithm requiring an RGB IHC image as input^41^ and outputting a nuclear mask based on the hematoxylin expression of cells. STARDIST requires the training model metadata for its segmentation, which we have added to MARQO’s docker environment. To reconcile multiple segmentation masks per batch of tiles, MARQO iterates across each identified nuclear object. If the centroid of that nuclear object is identified across at least several iterative stains (by default 60%) within a set distance (by default 3 µm) to account for any registration error margins (both tunable parameters), then that nuclear object is kept in the final composite segmentation mask. Reconciling across multiple stains permits the pipeline to have a larger sample size to evaluate whether a cell is a true positive-segmented cell versus an RBC, an artifact, or a cell lost from tissue damage that was deemed a false positive-segmented cell. For technologies with only one nuclear stain, MARQO performs a singular segmentation and uses this nuclear mask for downstream quantification. For singleplex IHC using a hematoxylin nuclear stain, STARDIST is similarly used. For mIF staining, a STARDIST module trained on a single channel nuclear stain 4′,6-diamidino-2-phenylindole (DAPI) is used, similarly outputting a semantic nuclear mask. Whether analyzing multiplex or singleplex data, the final cell segmentation mask ensures that each cell is only segmented only once.

#### MARQO quantification module

In its quantification module, MARQO imports the predetermined nuclear semantic mask and expands the nuclear boundaries by a predetermined number of pixels to simulate an artificial cytoplasm. The user defines how many pixels to expand each nucleus by per stain in the launch quality control. The module acquires pixel and morphological metadata for each cell’s nucleus, cytoplasm, and membrane. These data include signal intensity values for both the chromogen stain and counterstain (minimum, maximum, percentiles, standard deviation, etc.) and the perimeter, area, major and minor axes for each cell. The overall median nuclear staining intensity is also quantified, and from this value, we can hypothesize whether a cell is an RBC, since it would not have a nucleus and retain a nuclear counterstain. Because adjacent batches of tiles analyzed have overlapping borders, cell information is only collected from nuclei centroids within the bounds of the original tile that were not extended. Metadata from all batches of tiles are appended to the features table, which contains metadata for all registered and segmented cells for all stains for the sample being analyzed.

A “.geojson” file format is produced containing all spatial metadata for all the segmented nuclei and cytoplasms. If the user prefers to use MARQO only for the registration and segmentation features and relies on a third-party tool (such as QuPath) to classify their cell types with methods not offered by MARQO (such as a fully supervised approach like neural networks), they may import these files into other software packages along with the raw images.

#### MARQO classification modules

MARQO offers two classification options for users. Both options leverage unsupervised learning to cluster all segmented and identified cells from the sample being analyzed, producing a pre-defined number of unique cell clusters per marker. After either option, the user determines which clusters contain true-positive or true-negative cells in the review GUI. To aid this decision process, the application overlays cell centroids per cluster over the raw images and visualizes any tile index that the user wants to inspect. Additionally, the user may fine-select individual cells to be considered positive or negative. After all the markers’ clusters are evaluated by the user, the final summary table is updated to indicate which markers each cell is either positive or negative for. This step can be done per sample or per group of samples at once.

The first classification option performs principal component analysis (PCA) on the metadata features table, reducing the dimensionality of the features table. The results from the PCA that capture 90% of the variance are used to cluster the data into a predefined number of clusters using the mini batch K-Means algorithm. This quicker clustering approach allowed us to implement a “re-classify” option for determined clusters directly within the GUI. If the user deems a cluster inadequate as completely positive or negative for all the cells inside it, the user may “re-classify” a cluster to output a new round of sub-tiers within the original tier. Because of the reduced dimensionality and quicker analysis, we recommend the k-means approach for larger samples and samples warranting quick re-clustering.

The second classification option iterates through every random 50,000 cells in the features table, which then undergo a PCA analysis, followed by Uniform Manifold Approximation and Projection (UMAP). The produced projection is clustered using a Gaussian Mixed Model (GMM), initiated by k centroids around which other centroids are clustered. This process produces approximately 60 GMMs across different batches for the same marker, which are then collapsed into the pre-defined number of clusters by standard K-means clustering of GMM means for each z-transformed feature.

### Statistical analysis

Statistical analyses were performed using Prism V.10 (GraphPad, California, USA). Multigroup analyses of variance were performed using one-way analysis of variance (ANOVA) followed by Šídák one-way comparison tests. The random model of precision versus recall represents the performance of a classifier that makes random guesses, achieving an area-under-the-curve (AUC) of 0.5. Non-parametric Spearman’s correlation values and linear regression analyses were used when comparing MARQO with manual pathologist analyses on QuPath.

## DATA AVAILABILITY

Image files and data are available upon request from the corresponding author.

## CODE AVAILABILITY

A container with MARQO’s GUI to analyze samples locally is available at https://github.com/igorafsouza/MARQO. All source code is available from the corresponding author upon reasonable request. Requests for service to run larger samples on a cluster should be made by contacting the corresponding author.

### Human subjects

HCC liver samples used for MICSSS and singleplex IHC were obtained via a single-arm, open-label, phase 2 trial of HCC patients with resectable tumors (ClinicalTrials.gov, <u>NCT03916627</u>, Cohort B). Twenty patients were enrolled and received two cycles of cemiplimab prior to surgical resection, as described in the clinical trial publication, including the full protocol provided in the supplementary materials^25^. Core needle biopsies and tumor tissues were obtained from these patients undergoing surgical resection at Mount Sinai Hospital (New York, NY), after obtaining informed consent in accordance with a protocol reviewed and approved by the Institutional Review Board at the Icahn School of Medicine at Mount Sinai (IRB 18-00407). Tonsil samples used for singleplex IHC were obtained from patients undergoing tonsillectomies, from Leica Biosystems, which purchased FFPE blocks from the Deer Park local hospital. Non-small cell lung cancer (NSCLC) resection samples used for MICSSS and mIF were obtained from treatment-naive patients undergoing surgical resection at Mount Sinai Hospital (New York, NY) after obtaining consent in accordance with a protocol reviewed and approved by the Institutional Review Board at the Icahn School of Medicine at Mount Sinai (IRB 21-01308). NSCLC, head and neck squamous cell carcinoma (HNSCC), colorectal cancer (CRC), breast cancer (BC), epithelial ovarian cancer (EOC), pancreatic ductal adenocarcinoma (PDAC), glioblastoma (GBM), renal cell carcinoma (RCC), and melanoma (MEL) samples used for the TMA MICSSS analysis were obtained by the Cooperative Human Tissue Network in a deidentified fashion without the possibility of being linked with metadata.

### Sample preparation and staining for singleplex IHC, MICSSS, and tissue microarray

For MICSSS and singleplex IHC, pretreatment HCC core biopsies and post-treatment surgically resected HCC lesions were obtained from formalin-fixed and paraffin-embedded (FFPE) blocks. Tissues used to create the TMA were similarly processed, and 1.0 mm diameter punches were made. FFPE tissue sections were sliced at 4_µm. Whole slide tissue sections were stained either by singleplex IHC or by using the MICSSS protocol as previously described^5^. The TMA was stained with MICSSS. All slides went through an automated immunostainer (Leica Bond RX, Leica Biosystems) which performed baking, chromogenic revelation, and nuclear counterstaining. Among the singleplex IHC, tonsil samples were subjected to chromogenic revelation via 3,3′-Diaminobenzidine (DAB, Vector Laboratories) and tumor samples, as well as all MICSSS samples underwent chromogenic revelation via 3-amino-9-ethylcarbazole (AEC, Vector Laboratories). All IHC assays were counterstained with hematoxylin. Then, all slides were mounted with a glycerol-based mounting medium and scanned to obtain digital images using an Aperio AT2 scanner and Aperio ImageScope DX visualizer software v.12.3.3 (Leica). For MICSSS, after each round of staining and scanning, slide coverslips were removed in hot water (∼50_°C) and tissue sections were bleached. This process was repeated for the length of the panel. Primary antibodies are presented in Supplementary Table 2 for the singleplex IHC, TMA, whole-slide resections and biopsies.

### Multiplex immunofluorescence staining and imaging on COMET^TM^

COMET mIF, or automated hyperplex immunofluorescence (IF), staining and imaging was performed on the COMET™ platform (Lunaphore Technologies). Slides from FFPE blocks were cut at 4µm and underwent 10 cycles of iterative staining and imaging, followed by an elution of the primary and secondary antibodies^9,42^. In brief, slides were preprocessed with PT Module (Epredia) with Dewax and HIER Buffer H (TA999-DHBH, Epredia) for 60min at 102°C. Subsequently, slides were rinsed and stored in Multistaining Buffer (BU06, Lunaphore Technologies) until use. The 20-plex protocol template was generated using the COMET™ Control Software, and reagents were loaded onto the device to perform the sequential immunofluorescence (seqIF™) protocol. A list of primary antibodies with corresponding dilution and incubation times is enclosed in Supplementary Table 3. Secondary antibodies were used as a mix of 2 species-complementary antibodies. The nuclear signal was detected with DAPI (Thermo Scientific, cat no: 62248, 1/1000 dilution) after 2 min of dynamic incubation. All reagents were diluted in Multistaining Buffer (BU06, Lunaphore Technologies). For each cycle, the following exposure times were used: 50 ms for DAPI, 400 ms for TRITC, and 200 ms for Cy5. The elution step lasted 2 min for each cycle and was performed with elution buffer (BU07-L, Lunaphore Technologies) at 37°C. The quenching step lasted for 30 sec and was performed with quenching buffer (BU08-L, Lunaphore Technologies). The imaging step was performed with imaging buffer (BU09, Lunaphore Technologies). The seqIF™ protocol in COMET™ resulted in a multi-layer “.ome.tiff” file where the imaging outputs from each cycle were stitched and aligned. COMET™ ome.tiff contains a DAPI image, intrinsic tissue autofluorescence in the TRITC and Cy5 channels, a single fluorescent layer per marker, and a single layer per additional image post-elution. Antibody titration was optimized to identify the best antibody dilution and incubation time. Imaging was performed on unstained tissue after each cycle of biomarker staining and antibody elution. The images were used to assess the staining quality and the elution efficiency for each staining condition.

### Image analysis performed on QuPath

To assess the performance of MARQO, we analyzed tiles from images produced by singleplex IHC, MICSSS, and COMET mIF on a third-party analysis tool commonly used in pathology labs, QuPath^19^. For the images produced from singleplex IHC and for each separate image produced from MICSSS, color deconvolution was performed for proper channel separation of the different stain vectors: hematoxylin, AEC or DAB chromogens, and the residual channel. For each whole slide image, these stain vectors were estimated from a selected sample ROI that included a balanced level of positive and negative cells as well as a blank white area to avoid image downsizing, which could affect the quality of the deconvolution. For the 500×500µm ROIs that were selected by a pathologist for MARQO segmentation and classification validation, stain vectors were estimated from the entire area of the ROI since QuPath does not downsize images this small. This deconvolution step was skipped for the “.ome.tiff” images produced by COMET mIF since each individual stain already existed in separate channels. Then, a whole tissue annotation was created using the “simple tissue detection” feature to automatically identify the whole tissue from the background glass. For the TMA images, separate tissue annotations were made for each individual core. Individual cells were then identified using STARDIST, a pre-trained nuclear segmentation algorithm that outputs a nuclear mask based on the hematoxylin or DAPI expression of the cells and includes cytoplasmic and membranous portions by expanding the nuclear segmentation border by a selected diameter. Hematoxylin and chromogen color intensity values for nuclear, cytoplasmic, and total cellular compartments as well as morphological features were recorded for each cellular detection. For each separate biomarker and each separate dataset, a machine learning classifier was trained by processing the hematoxylin/chromogen intensity and morphological values from a proper number of manually selected positive and negative cells in the cohort. The classifier was then applied to all the images stained for that biomarker in the entire cohort to identify positively stained cells. Given the heterogeneous nature of the different tissue types included in the TMA, applying these high-level classifiers for individual biomarkers across all cores regardless of tissue type was not always sufficient for proper identification of positively stained cells. When this was the case, more focused classifiers, that were trained on and applied to individual biomarkers within only one tissue type, were used for improved identification of positively stained cells. Finally, cell marker density data (number of positive cells/total tissue area) were exported^7^.

### Imaging proximity analysis

Using the features table, we extracted the centroid coordinates of all cells and stratified them based on their marker combination patterns and localization within the annotated tissues. For each cell in population A, we computed the distances to the nearest neighbor in population B. Based on the collected distribution of minimal distances, we computed a Gaussian kernel density estimation (KDE) curve and estimated the mode. We repeated this process for all possible pairs between cell types A and B for each patient-sample stratified per region of interest. Finally, we plotted the distribution of modes from the CD8 T cell perspective to themselves, B cells, and Tregs across tissue regions. This analysis will be available as a review module in a future deployment of the pipeline.

## SUPPLEMENTARY TABLES

**Supplementary Table 1:**
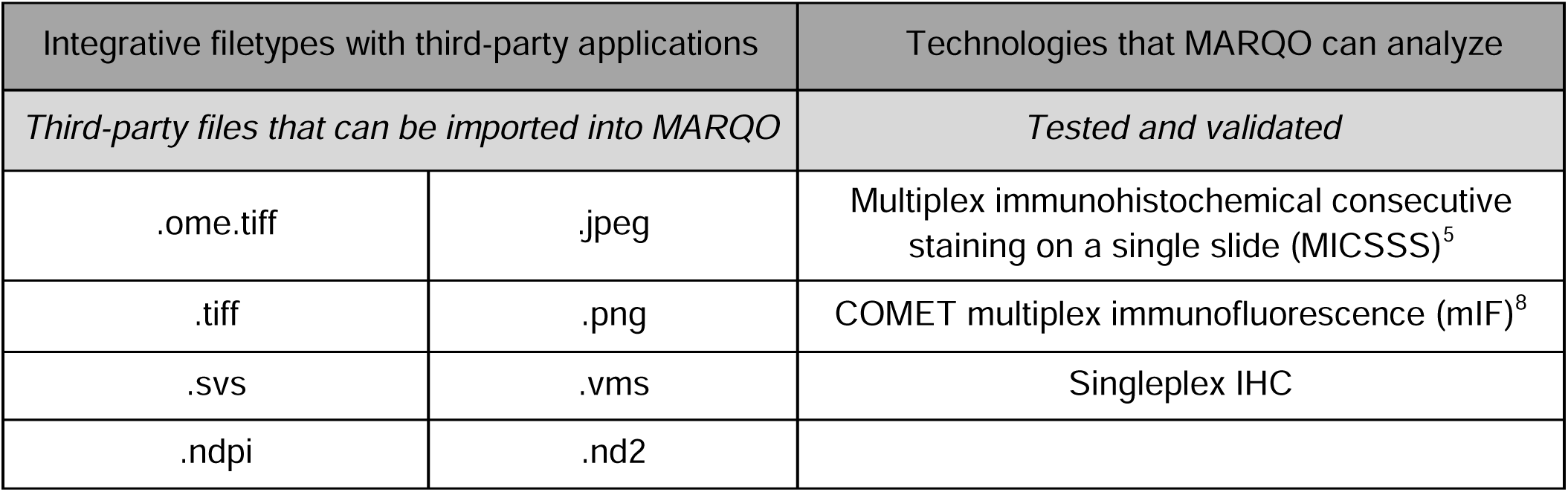
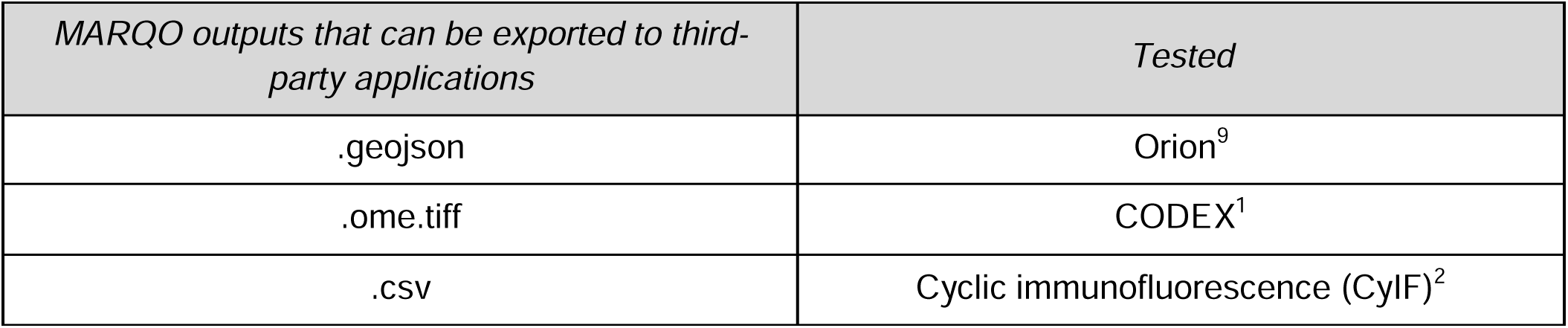
MARQO compatibility to import and export files with third-party applications (left) and its validation on multiple staining technologies (right)

**Supplementary Table 2:** Primary antibodies used for singleplex IHC, MICSSS on TMA, and MICSSS on biopsy and whole-slide HCC and NSCLC resections

| Cycle | Marker | Detection chromogen | Antibody clone | Antibody supplier | Antibody Dilution | Antibody incubation time (min) | Chromogen incubation time (min) |
| --- | --- | --- | --- | --- | --- | --- | --- |
| *Singleplex IHC* |  |  |  |  |  |  |  |
| 1 | CD3 | DAB | LN10 | Leica Biosystems | RTU | 15 | 10 |
| 1 | PD-L1 | AEC | E1L3N | Cell Signaling Technology | 1/400 | 15 | 15 |
| *MICSSS on TMA* |  |  |  |  |  |  |  |
| 1 | PD-L1 | AEC | E1L3N | Cell Signaling Technology | 1/400 | 15 | 15 |
| 2 | PD-1 | AEC | EPR4877(2) | Abcam | 1/250 | 15 | 15 |
| 3 | CD8 | AEC | C8/144b | Dako | 1/100 | 15 | 3 |
| 4 | CD3 | AEC | 2GV6 | Ventana | RTU | 15 | 3 |
| 5 | PanCK | AEC | AE1/AE3 | Dako | 1/50 | 15 | 15 |
| 6 | FoxP3 | AEC | 236A/E7 | Abcam | 1/80 | 15 | 15 |
| *MICSSS on biopsy and whole-slide HCC resections* |  |  |  |  |  |  |  |
| 1 | PD-1 | AEC | EPR4877(2) | Abcam | 1/250 | 15 | 15 |
| 2 | FoxP3 | AEC | 236A/E7 | Abcam | 1/80 | 15 | 15 |
| 3 | CD3 | AEC | 2GV6 | Ventana | RTU | 15 | 15 |
| 4 | CD8 | AEC | C8/144b | Dako | 1/100 | 15 | 15 |
| 5 | Ki-67 | AEC | 30-9 | Ventana | RTU | 15 | 15 |
| 6 | αSMA | AEC | 1A4 | Dako | 1/250 | 15 | 15 |
| 7 | CD68 | AEC | KP1 | Dako | 1/1000 | 15 | 15 |
| 8 | CD20 | AEC | L26 | Dako | 1/250 | 15 | 15 |
| 9 | PanCK | AEC | AE1/AE3 | Dako | 1/50 | 15 | 15 |
| 10 | MZB1 | AEC | Polyclonal | Sigma Aldrich | 1/5000 | 15 | 15 |
| MICSSS on whole-slide NSCLC resections |  |  |  |  |  |  |  |
| 1 | PD-L1 | AEC | E1L3N | Cell Signaling Technology | 1/400 | 15 | 15 |
| 2 | FoxP3 | AEC | 236A/E7 | Abcam | 1/80 | 15 | 15 |
| 3 | PD1 | AEC | EPR4877(2) | Abcam | 1/250 | 15 | 15 |
| 4 | αSMA | AEC | 1A4 | Dako | 1/250 | 15 | 5 |
| 5 | CD68 | AEC | KP1 | Dako | 1/1000 | 15 | 5 |
| 6 | Ki-67 | AEC | 30-9 | Ventana | RTU | 15 | 3 |
| 7 | CD8 | AEC | C8/144b | Dako | 1/100 | 15 | 3 |
| 8 | CD3 | AEC | 2GV6 | Ventana | RTU | 15 | 3 |
| 9 | CD20 | AEC | L26 | Dako | 1/250 | 15 | 5 |
| 10 | PanCK | AEC | AE1/AE3 | Dako | 1/50 | 15 | 15 |

\*RTU = Ready-to-use

**Supplementary Table 3:**
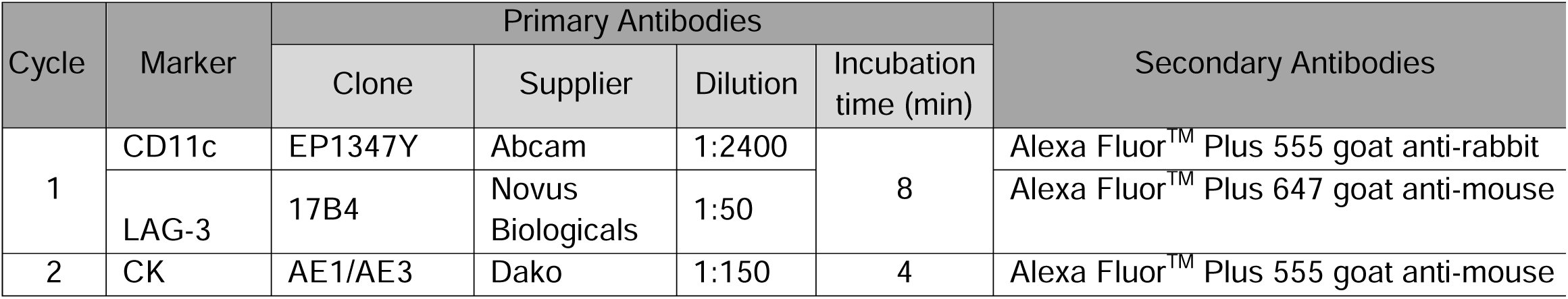
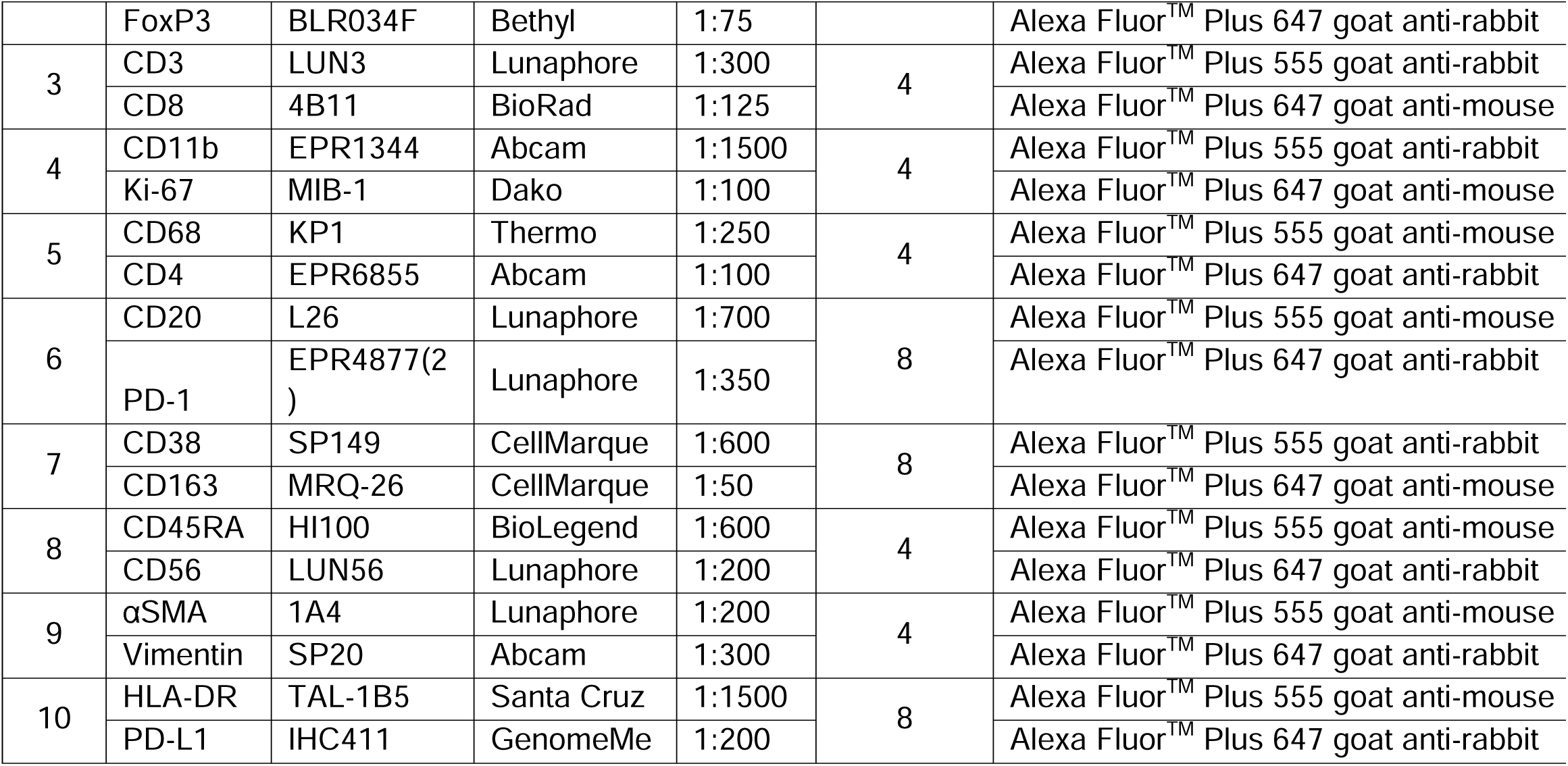
Primary and secondary antibodies used for the 20-plex seqIF™ protocol on COMET™ on NSCLC resections

## ACKNOWLEDGEMENTS

We thank the patients and their families for participating in our research study and for providing clinical trial and donor specimens. We thank the Biorepository and Pathology CoRE Laboratory of the Icahn School of Medicine at Mount Sinai for support. We also thank members of the Merad and Gnjatic laboratories for their help and support. This work was supported in part through the computational and data resources and staff expertise provided by Scientific Computing and Data at the Icahn School of Medicine at Mount Sinai and supported by the Clinical and Translational Science Awards (CTSA) grant UL1TR004419 from the National Center for Advancing Translational Sciences. The clinical trial (ClinicalTrials.gov, <u>NCT03916627</u>, Cohort B) and part of this project were funded by Regeneron Inc. S.G. was partially supported by National Institutes of Health (NIH) grants CA224319, DK124165, CA263705 and CA196521. M.M. was partially supported by NIH grants CA257195, CA254104 and CA154947. T.U.M. was partially supported by the Tisch Cancer Institute Cancer Center Support Grant (P30 CA196521).

## AUTHOR CONTRIBUTION

M.B., P.H., S.T., S.H., M.M., and S.G. conceptualized the project. T.U.M., M.M., and S.G. obtained funding for the project. M.B., G.I., P.H., and S.G. designed the experiments. M.B., I.F., G.I., V.R., E.G-K., S.O., R.C., L.T., J.LB., S.H., and G.A. performed the experiments. M.B., I.F., G.I., P.H., and S.G. analyzed the experiments. M.S. and T.U.M. provided all clinical care to patients in the clinical trial. S.O., R.C., Z.Z., S.C.W, M.I.F., and R.B. provided expertise for pathological response assessment and tissue annotation. C.H. coordinated the clinical and research teams. M.B. and I.F. developed the pipeline. M.B., I.F., and V.R. performed computational analysis. S.T., E.G-K., S.K-S., M.M., and S.G. provided intellectual input. M.B. wrote the manuscript. P.H. and S.G. edited the manuscript. All authors provided feedback on the manuscript draft.

