## Supplemental Figures for "MARQO pipeline resolves multiparametric cellular and spatial organization in cancer tissue lesions"

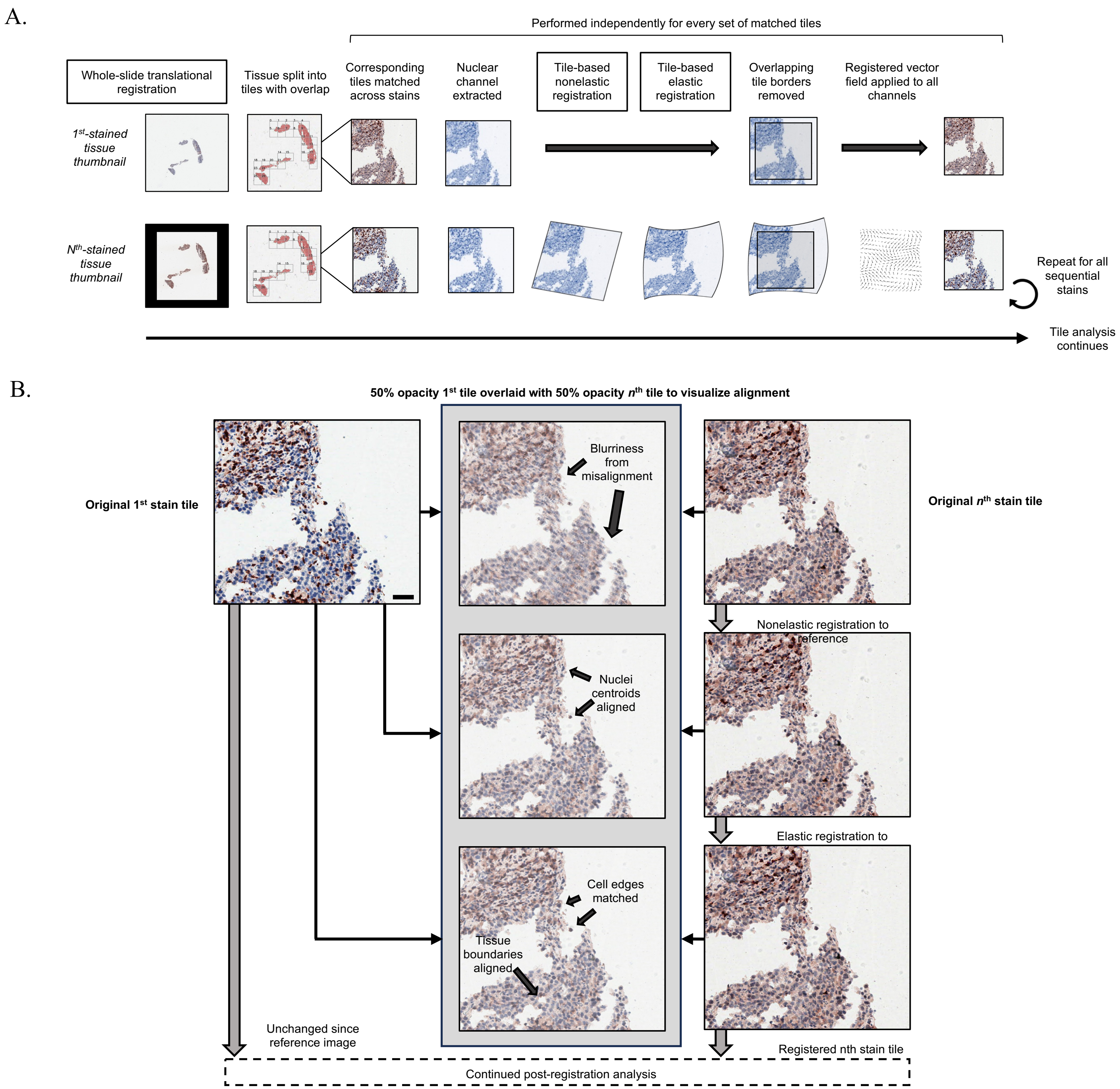


**Figure S1: Rigid and elastic registrations methodology with examples. A)** MARQO performs an alignment methodology consisting of three distinct registration steps. At first, during the initial user quality control MARQO performs a translational registration of low-resolution thumbnail images, aligning each sequential stain to the first stain thumbnail, which the user can deem as appropriate or revise with tunable, intuitive parameters in the interactive application (see Supplementary Video 1). After tiling of all sequential high-resolution images, MARQO matches corresponding tiles, extracts the nuclear stain, and performs both rigid affine and elastic "B-Spline" registrations on this channel. Tiles were analyzed with overlap which are then removed. The registration vector field from the nuclear channel is applied to the original red-green-blue (RGB) tile so that all tiles are now registered. This process repeats for all tiles and stains, which by default is done in parallel. Analysis per batch of tiles then continues through the MARQO pipeline. **B)** Top row: An example 1st-stained tile (left) and its corresponding nth-stained tile (right) are shown after the initial thumbnail translational registration and tiling steps with an image depicting 50% transparency 1st-stained tile overlaid to 50% transparency nth-stained tile (middle). Note the severe misalignment at the cellular level with dark arrows. Middle row: the nth-stained tile after nonelastic, affine registration (right) and its overlay with the reference tile (middle). Note the aligned cell centroids with some minor misalignment at cell and tissue boundaries with dark arrows. Bottom row: the nth-stained tile after elastic, B-Spline registration (right) and its overlay with the reference tile (middle). Note enhanced cell and tissue boundary alignment with dark arrows. Scale bar = 50µm.

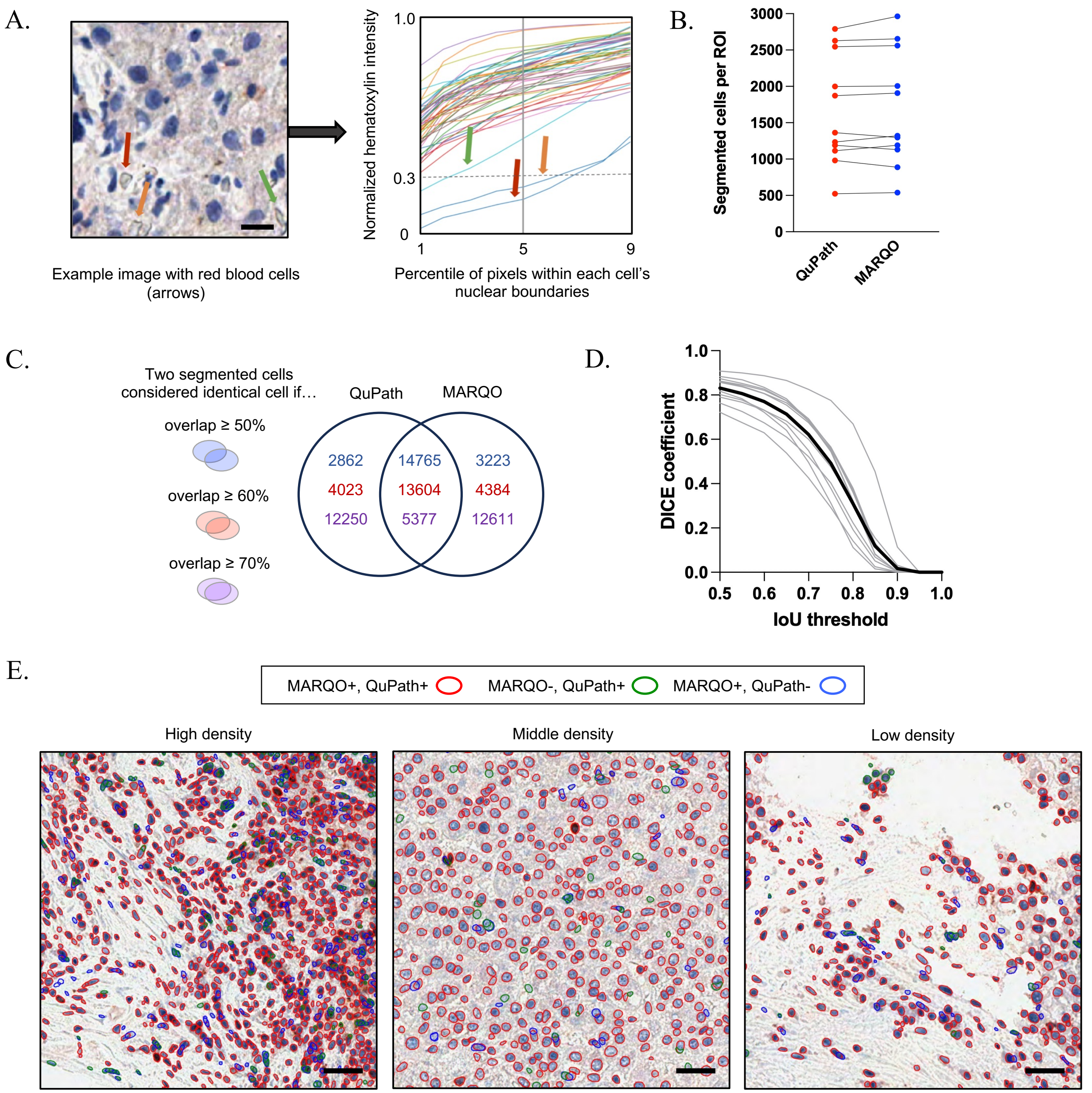


**Figure S2: MARQO segmentation performance. A)** Example image depicting red blood cells, shown by arrows. Adjacent graph plots the hematoxylin nuclear counterstain intensity for each cell along all its pixels' percentiles. Scale bar = 10μm. **B)** Bar graph depicting total number of cells segmented by a pathologist using QuPath versus MARQO's automatic composite segmentation for 10 tiles that spanned smaller biopsies and larger whole-slide tissue resections with heterogeneous cell densities. **C)** Comparing MARQO to QuPath nuclear segmentation, if a MARQO-segmented cell's boundaries and a QuPath-segmented cell's boundaries overlap with at least 50% area (blue), the Venn Diagram depicts the summation of all 10 tiles' cells segmented uniquely by MARQO, uniquely by QuPath, and by both. This is repeated for a 60% overlap threshold (red) and a 70% overlap threshold (purple). **D)** Similar to C, line graph plots the DICE coefficient across a range of Intersection over Union values (IoU), which are synonymous to overlap threshold divided by 100. Each gray curve represents one tile, while the thicker, black curve is the average of all tiles. **E)** Example tiles depict the performance of MARQO compared to QuPath across different cellular densities in whole-slide resected HCC tissue. Using an overlap threshold of 50%, or an IoU of 0.5, red signifies cells segmented by both MARQO and QuPath, green signifies cells segmented by QuPath but not MARQO, and blue signifies cells segmented by MARQO but not QuPath. Scale bars = 30μm.

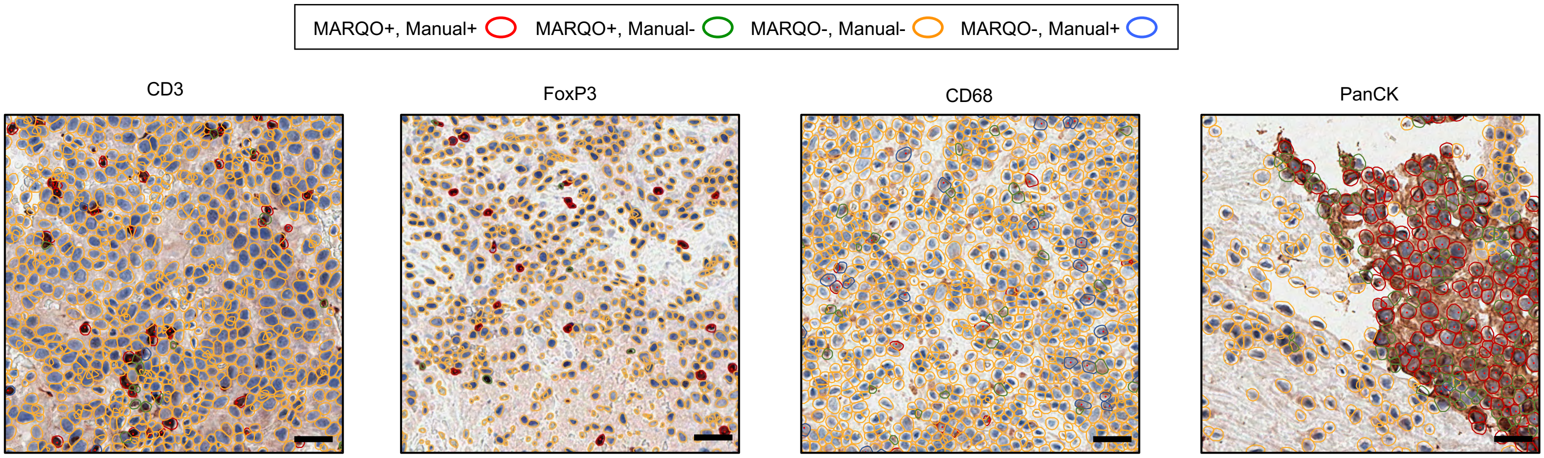


**Figure S3: Example tiles used for classification validation.** These cropped tiles correspond to the stacked bar graphs provided in Figure 3E across the markers CD3, FoxP3, CD68, and PanCK chosen for classification validation with the pathologist. Red signifies cells deemed positive by both MARQO after user quality control and the pathologist. Green signifies cells considered positive by MARQO after user quality control but not the pathologist. Blue signifies cells considered positive by the pathologist but not MARQO after user quality control. Yellow signifies cells considered not positive by both MARQO after user quality control and the pathologist. Scale bars = 30μm.

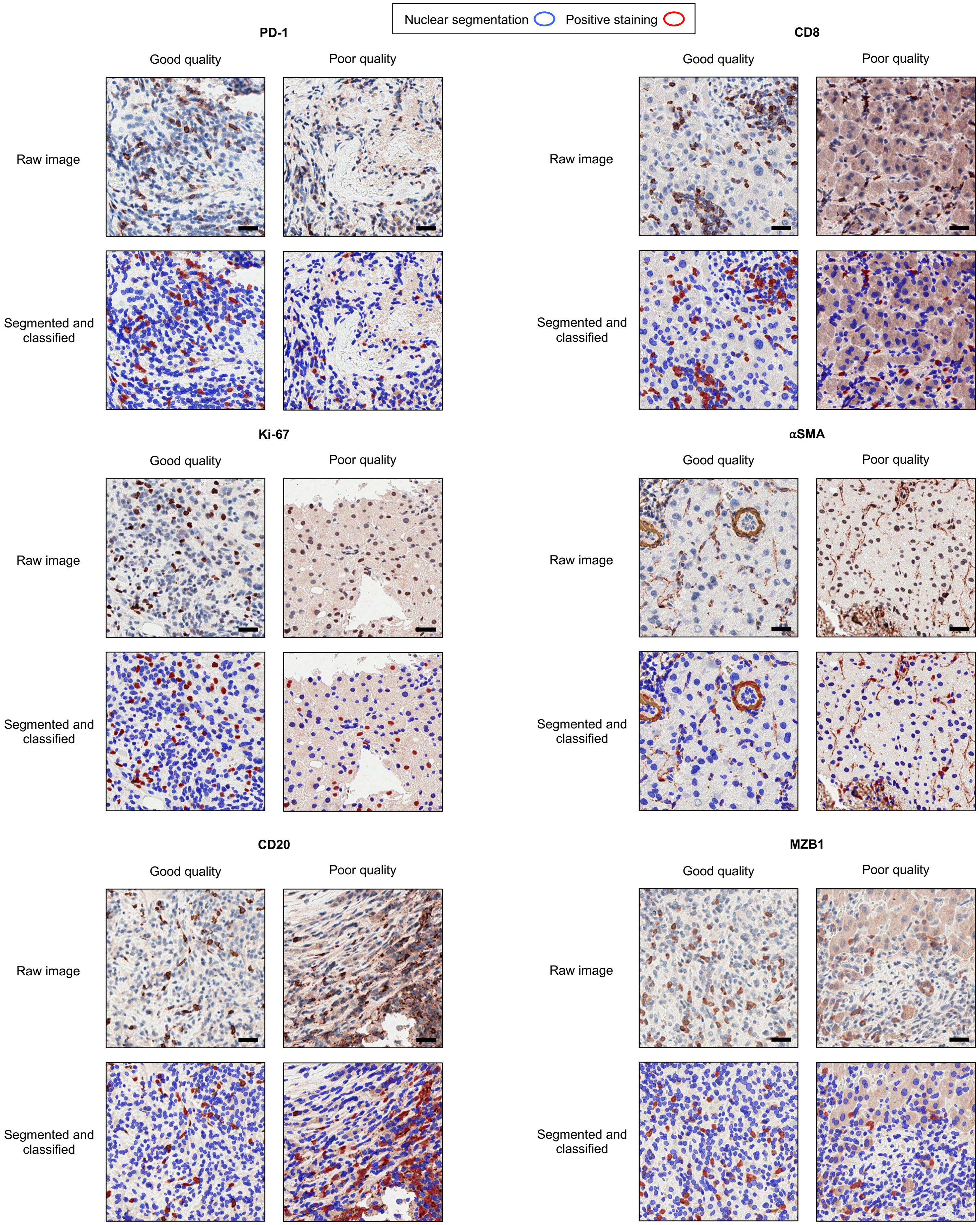


**Figure S4: Examples of good- and poor-quality tiles used to test MARQO.** These cropped tiles across diverse markers, including PD1, FoxP3, CD3, CD8, Ki-67, αSMA, CD68, NKp46, CD20, and PanCK, were segmented and classified by MARQO and the user with quality control. A pathologist ranked these tiles as “good” (left) and “poor” (right) qualities, defined based on staining, tissue damage, blurriness, or other factors. Scale bars = 30μm.
